# Association of Structural Variation with Cardiometabolic Traits in Finns

**DOI:** 10.1101/2020.12.13.422502

**Authors:** Lei Chen, Haley J. Abel, Indraniel Das, David E. Larson, Liron Ganel, Krishna L. Kanchi, Allison A. Regier, Erica P. Young, Chul Joo Kang, Alexandra J Scott, Colby Chiang, Xinxin Wang, Shuangjia Lu, Ryan Christ, Susan K. Service, Charleston W.K. Chiang, Aki S. Havulinna, Johanna Kuusisto, Michael Boehnke, Markku Laakso, Aarno Palotie, Samuli Ripatti, Nelson B. Freimer, Adam E. Locke, Nathan O. Stitziel, Ira M. Hall

**Affiliations:** McDonnell Genome Institute, Washington University School of Medicine, St. Louis, MO, USA; Department of Medicine, Washington University School of Medicine, St. Louis, MO, USA; Department of Genetics, Yale University School of Medicine, New Haven, CT, USA; Department of Genetics, Washington University School of Medicine, St. Louis, MO, USA; Cardiovascular Division, Department of Medicine, Washington University School of Medicine, St. Louis, MO, USA; Center for Neurobehavioral Genetics, Jane and Terry Semel Institute for Neuroscience and Human Behavior, University of California Los Angeles, Los Angeles, CA, USA; Center for Genetic Epidemiology, Department of Preventive Medicine, Keck School of Medicine, University of Southern California, Los Angeles, CA, USA; Quantitative and Computational Biology Section, Department of Biological Sciences, University of Southern California, Los Angeles, CA, USA; Institute for Molecular Medicine Finland (FIMM), HiLIFE, University of Helsinki, Helsinki, Finland; Finnish Institute for Health and Welfare (THL), Helsinki, Finland; Institute of Clinical Medicine, Internal Medicine, University of Eastern Finland, Kuopio, Finland; Department of Medicine, Kuopio University Hospital, Kuopio, Finland; Department of Biostatistics and Center for Statistical Genetics, University of Michigan School of Public Health, Ann Arbor, MI, USA; Analytical and Translational Genetics Unit (ATGU), Psychiatric & Neurodevelopmental Genetics Unit, Departments of Psychiatry and Neurology, Massachusetts General Hospital, Boston, MA, USA; Broad Institute of MIT and Harvard, Cambridge, MA, USA; Department of Public Health, Faculty of Medicine, University of Helsinki, Finland

## Abstract

The contribution of genome structural variation (SV) to quantitative traits associated with cardiometabolic diseases remains largely unknown. Here, we present the results of a study examining genetic association between SVs and cardiometabolic traits in the Finnish population. We used sensitive methods to identify and genotype 129,166 high-confidence SVs from deep whole genome sequencing (WGS) data of 4,848 individuals. We tested the 64,572 common and low frequency SVs for association with 116 quantitative traits, and tested candidate associations using exome sequencing and array genotype data from an additional 15,205 individuals. We discovered 31 genome-wide significant associations at 15 loci, including two novel loci at which SVs have strong phenotypic effects: (1) a deletion of the *ALB* gene promoter that is greatly enriched in the Finnish population and causes decreased serum albumin level in carriers (p=1.47×10^−54^), and is also associated with increased levels of total cholesterol (p=1.22×10^−28^) and 14 additional cholesterol-related traits, and (2) a multiallelic copy number variant (CNV) at *PDPR* that is strongly associated with pyruvate (p=4.81×10^−21^) and alanine (p=6.14×10^−12^) levels and resides within a structurally complex genomic region that has accumulated many rearrangements over evolutionary time. We also confirmed six previously reported associations, including five led by stronger signals in single nucleotide variants (SNVs), and one linking recurrent *HP* gene deletion and cholesterol levels (p=6.24×10^−10^), which was also found to be strongly associated with increased glycoprotein level (p=3.53×10^−35^). Our study confirms that integrating SVs in trait-mapping studies will expand our knowledge of genetic factors underlying disease risk.

## Introduction

Common human diseases affecting the cardiovascular and endocrine systems are known to be associated with a variety of quantitative risk factors including various measures of cholesterol, metabolites, insulin, glucose, blood pressure, and obesity. Understanding the genetic basis of these and other quantitative traits can shed light on the etiology, prevention, diagnosis, and treatment of disease. Family and population-based studies have shown significant heritability for many cardiometabolic traits, and prior genome-wide association studies (GWAS) have identified hundreds of associated loci. However, most prior trait-mapping studies have focused on common variants ascertained by genotyping arrays, or rare coding variants measured by exome sequencing, leaving out the contribution of larger and more complex forms of genome variation.

Of particular interest is the contribution of genome structural variation (SV), which encompasses diverse variant types larger than 50 base pairs (bp) in size, including copy number variants (CNVs), mobile element insertions (MEIs), inversions, and complex rearrangements. Although rare and *de novo* SVs have long been recognized to cause various rare and sporadic human disorders, and somatic SVs play a central role in cancer biology, the extent to which SVs contribute more generally to common diseases and other complex traits in humans is less clear. Early genome-wide studies^1–3^ failed to identify SVs associated with common diseases, but these were limited by the use of low-resolution array platforms, which only capture extremely large CNVs (>100kb, or similar), and by modest sample size. Several later studies performed targeted analysis of known SVs combined with larger-scale GWAS data^4–6^, leading to the association of structural alleles at *HP* and *LPA* with cholesterol levels. More recent array-based CNV association studies with large sample sizes (>50,000 individuals) have revealed several genome-wide significant CNV loci for anthropometric traits and coronary disease, but these studies focused on extremely large CNVs representing <1% of the overall SV burden, leaving most SVs untested^7–9^. Fine mapping of expression quantitative trait loci (eQTLs) using deep whole genome sequencing (WGS) data has indicated that SVs are the causal variant at 3.5-6.8% of eQTLs, and that causal SVs have larger effect sizes than causal single nucleotide variants (SNVs) and indels and are often not well-tagged by flanking SNVs^10,11^. This suggests that direct assessment of SVs in WGS-based complex trait association studies has the potential to reveal novel causative variants not found through other approaches.

Here, we have performed a SV association study using deep (>20x) WGS data from 4,030 individuals from Finland with extensive cardiometabolic trait measurements, and extended these results to a larger set of 15,205 individuals with whole exome sequencing (WES) and single nucleotide polymorphism (SNP) genotype data. Compared to prior work, our study benefits from (1) comprehensive SV ascertainment due to the use of deep WGS data and complementary SV detection methods, (2) deeply phenotyped individuals with existing SNP array and exome sequence data, and (3) the unique history of the Finnish population, which was shaped by multiple population bottlenecks and rapid population expansions, leading to an enrichment of some otherwise rare and low-frequency variants that can be detected by trait association at relatively modest sample sizes^12–14^. By testing for associations between structural variants and cardiometabolic traits, we identified 15 genome-wide significant loci, nine of which remained significant after multiple testing correction for the number of phenotypes, including a Finnish-enriched promoter deletion of the *ALB* gene associated with multiple traits, and a multiallelic CNV affecting the *PDPR* gene associated with pyruvate levels.

## Material and Methods

### Samples and phenotype collection

The genomic data in this study come from 10,197 METSIM participants collected from Kuopio in Eastern Finland, and 10,192 FINRISK participants collected from northeastern Finland. Both studies were approved by the Ethics Committees in Finland and all individuals contributing samples provided written informed consent. Besides collecting genotype data by SNP array and exome sequencing, both studies measured up to 254 quantitative cardiometabolic traits, among which we selected 116 traits with adequate sample sizes to maintain trait-mapping power (see below). All phenotype data were residualized for trait-specific covariates and transformed to a standard normal distribution by inverse normalization. Complete details of sample collection, genotype acquisition, and trait adjustments were described previously^14^.

### Power estimation and phenotype selection

Phenotypes with limited sample size are likely to be underpowered in trait-mapping analysis and increase the test burden if included. Thus, we selected 116 traits with large enough sample size that guaranteed 80% power to detect a hypothesized rare SV (Minor allele count (MAC) =10) with strong effect (explained 8.4% of the additive quantitative trait locus (QTL) variance, a contribution comparable to the effect of SV expression QTLs^10^). We estimated the minimum required sample size as 375 through an analytical approach implemented in Genetic Power Calculator^15^. Several other assumptions for the calculation are: 1. All samples are independent (sibship size=1); 2. The top signal is in perfect linkage disequilibrium (LD) with the causal variant; and 3. type I error rate=1×10^−6^.

### Generation of SV callsets from WGS data

For SV discovery, we used WGS data from 3,082 METSIM participants and 1,114 FINRISK participants sequenced at the McDonnell Genome Institute under the NHGRI Centers for Common Disease Genomics (CCDG) program. To increase variant detection sensitivity, we also included 779 additional Finnish participants from other cohorts and 112 multi-ethnic samples from 1000 Genomes (1KG) Project. All genomes were sequenced at >20x coverage on the Illumina HiSeq X and NovaSeq platforms with paired-end 150bp reads.

WGS data were aligned to the GRCh38 reference genome using BWA-MEM and processed using the functional equivalence pipeline^16^. An SV callset based on breakpoint mapping was generated using our recently published workflow^17^ using the same methods as in our recent study of 17,795 human genomes^18^. Briefly, we ran LUMPY (v0.2.13)^19^, CNVnator (v0.3.3)^20^, and svtyper (v0.1.4)^21^ to perform per-sample variant calling. After removing 22 samples that failed quality control, we merged sites discovered in all the samples and re-genotyped all sites in all samples to create a joint callset using svtools (v0.3.2)^17^. Each variant was characterized as either deletion (DEL), duplication (DUP), inversion (INV), mobile element insertion (MEI), or generic rearrangement of unknown architecture (BND), based on comprehensive review of its breakpoint genotype, breakpoint coordinates, genome annotation, and read-depth evidence, as described previously^17,18^. According to our definition of SV, we filtered variants smaller than 50bp. Moreover, we tuned the callset based on Mendelian error rate and flagged BNDs with mean sample quality (MSQ) score <250 and INVs with MSQ <100 as low-confidence variants. Details about this QC strategy are described elsewhere^18^. For convenience, we refer to this as the “LUMPY callset”.

We applied two read-depth based CNV detection methods to WGS data to detect variants that might be missed by breakpoint mapping. GenomeSTRiP^22^ is an established tool for cohort-level CNV discovery that has proven effective in many prior studies; however, when using the recommended parameters (as we did here), detection is limited to larger CNVs (>1kb) within relatively unique genomic regions. Thus, in parallel we used a custom cohort-level CNV detection pipeline based on CNVnator^20^ to detect smaller and more repetitive CNVs (see below).

We adapted the original GenomeSTRiP pipeline (v2.00.1774) for the large cohort of 5,087 Finnish samples: after the SVPreprocess step, samples were grouped by study cohorts and sorted by sequencing dates, then split into 54 batches with maximum size of 100. CNVs were detected within each batch by CNVDiscoveryPipeline and classified as either deletion (DEL), duplication (DUP), or mixed CNV (mCNV), with both copy number gain and loss existing in the population (referred to as “multiallelic CNV” in the text). Next, we concatenated variants from the 54 batch VCFs and re-genotyped all variants in all samples using SVGenotyper to produce a joint callset. Then we ran several GenomeSTRiP annotators (CopyNumberClassAnnotator, RedundancyAnnotator) to reclassify variants and remove redundant variant calls. During callset generation, 72 samples with abnormal read-depth profiles were excluded.

The read-depth based “CNVnator” callset was constructed using a custom pipeline that took as inputs the individual-level CNV callsets generated by CNVnator during the svtools pipeline. After removing samples with abnormal read-depth profiles, CNV calls from 4,979 samples were sorted and merged using the svtools pipeline. All merged CNV calls were re-genotyped in all samples using CNVnator. Within each connected component of overlapping CNV calls, individual variant calls were clustered based on correlation of copy-number profiles and by pairwise overlap. For each cluster, a single candidate was chosen to represent the underlying CNV. For sites with carrier frequency >0.1%, we fit the copy number distribution to a series of constrained Gaussian Mixture Models (GMMs) with varying numbers of components, and selected the site with the “best” variant representation based on a set of model metrics, including the Bayesian Information Criterion (BIC) and the distance between cluster means (“mean_sep”). For the remaining sites we selected those with the most significant copy number difference between carriers and non-carriers. With the same criteria used in GenomeSTRiP, we assigned integer copy number genotypes and CNV categories to the variants.

We used array intensity data for 2,685 METSIM samples to estimate the false discovery rate (FDR) under different filtering criteria, and to tune both CNV callsets. FDR was estimated from the Intensity Rank Sum (IRS) test statistics based on CNVs intersecting at least two SNP probes. Based on the FDR curves (**Figure S1**) we excluded GenomeSTRiP variants with GSCNQUAL score<2 and CNVnator DELs and DUPs with mean_sep < 0.47 or low carrier counts (DUPs<1, DELs<5, mCNVs<7).

To eliminate likely false positive calls introduced by sequencing artefacts, we excluded 612 LUMPY SVs, 740 GenomeSTRiP SVs, and 1098 CNVnator SVs that were highly enriched in any of the three sequencing year batches (P<10^−200^ from Fisher’s exact test). We further excluded 3 samples in the LUMPY callset, 72 samples in the GenomeSTRiP callset, and 12 samples in the CNVnator callset that carried abnormal numbers of variants (outlier samples defined by the difference of per-sample SV count from median divided by mad larger than 10 for LUMPY/GenomeSTRiP or larger than 5 for CNVnator). Together with the samples that failed QC during variant calling, the combined list of outliers consists of 84 METSIM samples, 56 FINRISK samples, and 99 samples from other cohorts. More information about sample- and variant-level exclusions can be found in **Table S1**.

For each high-confidence callset, we evaluated the final FDR by using the IRS, and ran the TagVariants annotator in GenomeSTRiP to estimate the proportion of SVs in LD with nearby SNPs (R_max_^2^>=0.5, flanking window size=1Mb). We calculated the overlap fraction between SV callsets by bedtools^23^ intersect (v2.23.0) requiring >50% reciprocal overlap between variants. To evaluate the genotype redundancy within and between callsets, we compared the original variant counts and the equivalent number of independent genetic variables estimated by a matrix decomposition method implemented in matSpDlite^24^, using the genotype correlation matrix as input. The space clustering was evaluated by running bedtools cluster with -d (max distance) specified as 10bp.

### Association test with WGS data

For CNV callsets, we defined minor allele count (MAC) as the number of samples with different genotypes from the mode copy number. We kept the conventional MAC definition for the LUMPY callset since it primarily contains biallelic SVs. We set the minimum MAC threshold as 10 for variants to be included in the trait association test. We renormalized the phenotype data of the WGS samples by rank-based inverse normal transformation. We performed single-variant association tests across all renormalized metabolic traits using the EMMAX model^25^ implemented in EPACTS (v3.2.9) software^26^. In the model, we specified the input genotype variables as the integer copy number genotype for GenomeSTRiP variants, allele balance for LUMPY variants, and raw decimal copy number for CNVnator variants. We also incorporated in the model a kinship matrix derived from SNP data by EPACTS to account for sample relatedness and population stratification.

We applied matSpDlite^24^ to estimate the equivalent number of independent tests. The genome-wide significance threshold was set at 1.89×10^−6^ after Bonferroni correction at level α = 0.05 over 26,495 independent genetic variables, and the experiment-wide significance threshold was set as 3.32×10^−8^ to further correct for the 57 independent phenotypic variables also estimated using matSpDlite^24^.

### Replication using exome and array data

We attempted to replicate the association signals with a nominal p<0.001 in WGS analysis using genotype data for an additional ∼15,000 FinMetSeq participants. To achieve this, we employed two approaches to infer the genotypes of candidate SVs from WES and array data: WES read depth analysis for CNVs and genotype imputation for biallelic SVs.

We separated the WES alignment data into two batches: the first composed of 10,379 samples sequenced with 100bp paired-end reads and the second composed of 9,937 samples sequenced with 125bp paired-end reads. For samples in each batch, we calculated the per-sample per-exon coverage by GATK^27^ DepthOfCoverage (v3.3-0) and adopted the data processing steps from the XHMM (v1.0) pipeline^28^ to convert the raw coverage data into PCA-normalized read-depth z-scores. Duplicated and outlier samples were filtered simultaneously, with 9,537 samples left in batch1 and 9,864 samples left in batch2. We calculated the correlation between SV genotypes from WGS data and the normalized read-depth z-scores of exons intersected or nearby (<5kb) using samples with both WES and WGS data. Exons with R^2^<0.1 were filtered out and the rest were passed on to validation, restricted to samples absent from the WGS analysis (n=15,205). The genetic relationship matrix used for WES replication was generated in a previous study^14^. We later did a meta-analysis under a fixed effect model using METASOFT (v2.0.1)^29^ to combine the results from the two WES batches, considering the two sequencing batches were actually sampled from the same population.

We converted the copy number genotypes (CN=2,3,4…) of 2,291 biallelic candidate SVs to allelic genotype format (GT=0/0, 0/1, 1/1) and extracted the SNPs and indels in the 1 Mb flanking regions of those SVs from the GATK callset generated from the same WGS data. We then phased the joint VCF with Beagle (version 5.1)^30^ to build a reference panel composed of 3,908 high-quality samples shared by the SV callset and the SNP callset. Then, we imputed the SV genotype in the additional 15,125 FinMetSeq samples with array genotype data by running Beagle on the genotyped SNPs. We filtered out low-imputation-quality SVs with DR2<0.3 reported by Beagle (the estimated correlation between imputed genotype and real genotype of each variant); then ran the EMMAX model on the 1,705 well-imputed SVs with the corresponding traits.

58 of the 2,053 candidate SVs had both imputed genotype and WES read-depth genotype, so we compared the imputation DR2 with exon-SV genotype R^2^, then chose the measurement better correlated with the WGS data. We then used Fisher’s method to combine the p-values from discovery stage (WGS data only) and replication stage. As a sanity check for the imputation quality, we conducted leave-one-out validation for the eight genome-wide significant SVs using the reference panel only. Specifically, we took one sample out each time as a test genome and imputed the SV genotype using the other 3,907 samples as reference and repeated the process 3,908 times to calculate the validation rate.

The array data and WES data were aligned to reference genome GRCh37 while the WGS data were aligned to reference genome GRCh38. For analysis, the coordinates were lifted over using the LiftOver utility from the UCSC GenomeBrowser (https://genome.ucsc.edu/cgi-bin/hgLiftOver).

### Candidate analysis

For genome-wide significant trait-SV associations, we collected previous GWAS signals on the same chromosome with P<10^−7^ from the EBI GWAS catalogue (https://www.ebi.ac.uk/gwas/docs/file-downloads, 2019-11-21 version) with the same set of keywords used in a previous study^14^ (one publication based on METSIM samples was excluded to only include findings from independent studies). We then performed conditional analysis on the original trait-SV pairs adding the GWAS hits as covariates. Conditional analyses were restricted to samples with WGS data to minimize the difference in genotype accuracy of the SV callset vs. the SNP callset.

For loci containing multiple genotype-correlated SVs associated with a trait, we lumped the variants together using bedtools merge^23^ and reported the coordinates of the entire region with the summary statistics of the strongest signal. To better understand these loci, we manually curated the candidates in IGV^31^ and extended the regions of interest to include surrounding genes, functional elements, previous GWAS signals and other genome annotations. We then equally split each region into ∼1000 windows and used CNVnator to calculate the copy number values of those windows for 100 individuals selected to represent all genotype groups. We then plotted the window-sample copy number matrix as a heatmap with scales best presenting the locus structure (e.g. **Figure 4**). In addition, for SNPs in the same region, we calculated the SNP-SV genotype correlation R^2^ by a linear regression model and SNP-trait p values by EMMAX, then plotted them together in a local Manhattan plot (e.g. **Figure 3**) using custom R scripts.

For the fine-mapping experiment of albumin, we selected the top 100 most significant SNPs on chr4:67443182-79382541 plus the *ALB* promoter deletion to calculate the pairwise genotype correlation matrix and ran CAVIAR (v0.2)^32^ on those 101 variants, with the “rho” probability set at 0.95 and varying the maximum number of causal variants one to five. The same experiment was done for total cholesterol. We used the model with maximum causal variants set at two to plot the posterior probability in **Figure 3**.

## Results

### Structural variation detection and genotyping

We identified 120,793 SVs by LUMPY^19^, 111,141 CNVs by GenomeSTRiP^22^ (GS), and 92,862 CNVs by our customized pipeline based on CNVnator^20^. Considering the different genotype metrics and detection resolutions, to retain sensitivity we chose to concatenate those three callsets together and adjust for redundancy later instead of merging the variants. 129,166 high-confidence autosomal SVs passed quality control, and 64,572 passed the frequency filter for association tests. **Figures 1** and **2** provide an overview of the high-confidence callset, including the composition, frequency, and size distribution broken down by SV types, biallelic vs. multi-allelic SVs, and detection pipelines. The SV size and frequency distributions are consistent with those in previous studies^10,18,33,34^: most called SVs are relatively small (<10kb), biallelic and rare; called MEIs exhibit the expected size distribution corresponding to Alu and L1 insertions; and allele frequency decreases with increased mean SV size, consistent with negative selection against large SVs (**Figure 2**).

**Figure 1.**
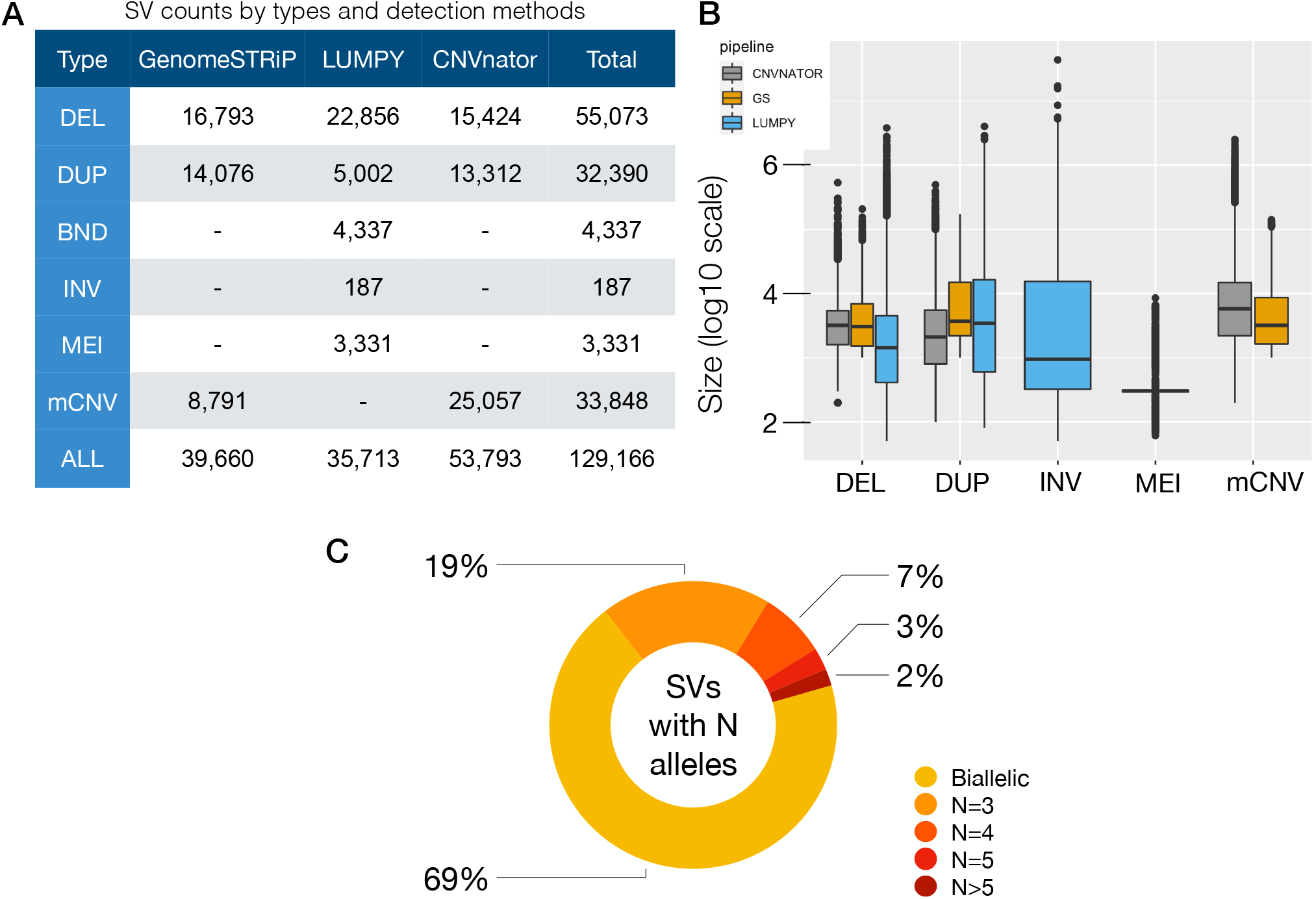
Overview of the high-confidence SV callset. **(A)** Count of high-confidence autosomal SVs stratified by variant type and detection method including deletions (DEL), duplications (DUP), multiallelic copy number variants (mCNV), inversions (INV), mobile element insertions (MEI) and generic rearrangements of unknown architecture (BND). **(B)** SV size distribution (log10 scale, bp) by variant type and detection method. mCNVs were only detected by read-depth based pipelines, INV and MEI variants were only detected in the LUMPY pipeline, and BNDs are not included due to the ambiguous definition of variant boundaries. **(C)** Proportion of bi-allelic SVs and multi-allelic CNVs, where N is defined by the number of copy number groups (e.g. CN=0,1,2,3,4, etc.)

**Figure 2.**
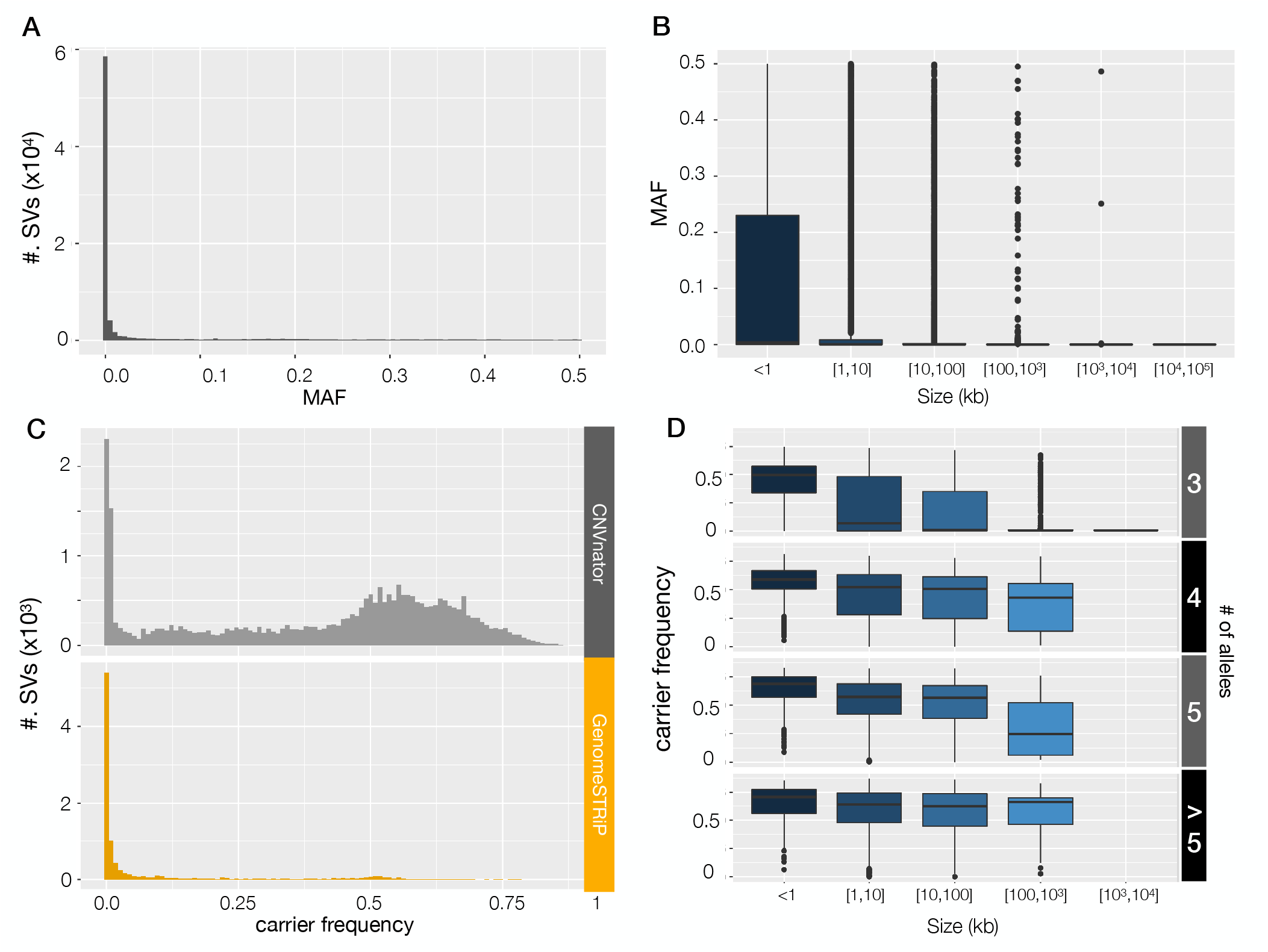
Frequency distribution of the high-confidence SVs. **(A)** The minor allele frequency distribution of all the high-confidence bi-allelic SVs and **(B)** the MAF distribution stratified by variant sizes. **(C) and (D)** are similar plots for multi-allelic CNVs, showing the frequency of carriers (non-diploid samples), stratified by detection methods. Note that the concentration of CNVnator variants between 0.5-0.75 were primarily caused by large segmental duplication regions near centromeres and telomeres, where the variant boundaries were challenging to define and the CNVs were detected in highly fragmented form. Such regions are often excluded from genetic analysis but were included here to maximize sensitivity.

Based on comparison with a set of SNP array intensity data (see **Methods**), we estimate an overall false discovery rate (FDR) of 4.7% for the high-confidence callset. As an indicator of true positive rate, the proportion of SV calls tagged by nearby SNPs (R^2^>=0.5, see **Methods**) was 56.8%, consistent with our prior GTEx study that used similar methods^10^ and was evaluated extensively in the context of eQTL mapping. We also compared our callset to the high-quality SV callsets from 1000 Genomes (1KG) and gnomAD projects and found an overlap of 35.2%, which is reasonable considering that these studies used distinct methods and sample sets. **Table 1** shows the above metrics stratified by pipelines. We estimated the genotype redundancy in total and stratified by pipelines (**Table S2**). Overall, the “effective sample size” of independent genetic variables was 55.5% of the original variant count. Additionally, since read-depth detection methods commonly result in “fragmented” CNV calls, we estimated the fragmentation level of calls by clustering variants within 10bp and measured the size of the clusters (**Table S3**).

**Table 1.** Callsets QC metrics Quality control metrics of the SV callsets including all variants, high-confidence variants, and high-confidence common variants (defined by >=10 carriers). CNV FDR was estimated by intensity rank sum test (IRS) using the SNP array data from METSIM samples. Note that LUMPY CNVs are by definition high confidence due to confirmation of independent read-depth support during variant classification steps (see **Methods**). Variant overlaps with 1KG and gnomAD were defined based on >50% reciprocal overlap. “Tagged by SNPs” was defined as SVs that are in LD (max r^2^>=0.5) with any SNP in the 1Mb flanking regions.

| QC Metrics | Variants Subset | LUMPY | GS | CNVNATOR |
| --- | --- | --- | --- | --- |
| CNV FDR* | all | - | 27% | 25% |
|  | high confidence | 0.80% | 3% | 9% |
| Counts | all | 120,793 | 111,141 | 92,862 |
|  | high confidence | 35,713 | 39,660 | 53,793 |
|  | common | 11,633 | 11,062 | 41,877 |
| Overlap w. 1kg* | all | 10% | 10% | 11% |
|  | high confidence | 34% | 21% | 15% |
|  | common | 49% | 34% | 13% |
| Overlap w. gnomad* | all | 18% | 14% | 25% |
|  | high confidence | 47% | 27% | 27% |
|  | common | 60% | 40% | 27% |
| Tagged by SNPs | high confidence | 63% | 62% | 46% |
|  | common | 77% | 65% | 49% |

\*CNVs only

Our CNVnator pipeline was the major source of redundancy and fragmentation since it detects CNVs with higher resolution – as small as 100bp – and covers repetitive and low-complexity regions, where the coverage profile is in general much noisier than the rest of the genome. The benefit is that CNVnator detected many true CNVs missed by the two other methods. As a benchmark of the sensitivity gain, we calculated the external validation rates for SVs uniquely detected in each of our pipelines (**Figure S2**). 7,210 variants identified only in CNVnator overlapped with variants in 1KG and gnomAD, contributing to the 43.1% of the overall CNVnator SVs that were validated through comparison to external datasets.

### Association of SVs with cardiometabolic traits

We first performed single variant association tests for 64,572 high-confidence SVs (MAC≥10) and 116 quantitative traits using the EMMAX model ^25^ in the 4,030 individuals with WGS data. We defined the genome-wide significance threshold as 1.89×10^−6^ and the experiment-wide significance threshold as 3.32×10^−8^ (see **Methods)**. Nine associations of six loci passed genome-wide significance threshold (**Table S5**); six were still significant after adjusting for the equivalent number of independent phenotypes (**Table 2**, WGS P).

**Table 2.** Summary statistics for all the genome-wide significant signals Summary statistics for 15 genome-wide significant loci with the top associated traits. Highly correlated SVs showing the same signal were manually inspected and clumped together. The genome-wide significance threshold was 1.89×10^−6^ and the experiment-wide significance threshold was 3.32×10^−8^ (see **Table S2** and **Methods** for details). The p value from WGS analysis and the p value from the replication experiment (IMP-imputation, WES-WES read-depth analysis, if applicable) were combined by Fisher’s method and used to determine the significance level. The carrier frequency was calculated in the WGS dataset. The column of “P GWAS conditioned” shows the SV p value conditioned on all intrachromosomal GWAS SNPs from GWAS Catalog^39^, using WGS data only (see **Methods**)

| SV type | Gene or annotation | Top trait | Chr | P WGS | P GWAS conditioned | BETA WGS | P combined | REP | Novel | Carrier frequency |
| --- | --- | --- | --- | --- | --- | --- | --- | --- | --- | --- |
| deletion | ALB | Albumin | 4 | 3.49E-21 | 1.05E-10 | 0.9107 | 1.47E-54* | IMP | Y | 0.03 |
| deletion | HP | Glyco-protein | 16 | 1.38E-10 | 3.63E-04 | -<br>0.1628 | 3.53E-35* | IMP | N | 0.55 |
| mCNV | PDPR | Pyruvate | 16 | 9.41E-11 | 1.07E-10 | -<br>0.7175 | 4.81E-21* | WES | Y | 0.02 |
| TCR | TRAV | CRP | 14 | 1.30E-15 | 1.89E-15 | 1.207 | 1.51E-16* | WES | Y | 0.36 |
| deletion | HNF1A-AS | CRP | 12 | 7.23E-04 | 3.60E-01 | 0.1912 | 4E-13* | IMP | N | 0.55 |
| TCR | TRBV | CRP | 7 | 3.36E-09 | 6.29E-09 | 0.8429 | 2.47E-16* | WES | Y | 0.38 |
| mCNV | NUMTS | Fast insulin | 1 | 1.00E-10 | NA | -<br>0.1179 | 1E-10* | NA | Y | 0 |
| MEI | LEPR | CRP | 1 | 3.94E-04 | 2.20E-01 | 0.164 | 4.5E-13* | IMP | N | 0.51 |
| deletion | IL34 | Tyrosine | 16 | 2.10E-04 | 5.45E-04 | 1.954 | 4.17E-10* | IMP | Y | 0.02 |
| MEI | CDH13 | Adiponectin | 16 | 1.24E-04 | 1.91E-02 | -<br>0.3282 | 3.68E-08 | IMP | N | 0.24 |
| mCNV | AMDHD1 | Histidine | 12 | 4.74E-04 | 2.72E-01 | 0.1485 | 5.33E-07 | IMP | N | 0.52 |
| mCNV | SegDup cluster | Fatty acid | 16 | 1.10E-06 | NA | -<br>0.1615 | 1.10E-06 | NA | Y | 0.57 |
| mCNV | SegDup cluster | Glutamine | 9 | 1.25E-06 | NA | -<br>0.7937 | 1.25E-06 | NA | Y | 0.43 |
| deletion | PLTP | Small HDL Particle | 20 | 2.40E-04 | 3.81E-02 | 0.1122 | 1.24E-06 | IMP | N | 0.53 |
| mCNV | Simple repeats | Creatinine | 4 | 1.41E-06 | NA | -<br>0.3949 | 1.41E-06 | NA | Y | 0.01 |

\* experiment-wide significant

We next sought to replicate these findings and to follow up on 4,855 loci with sub-threshold associations (p<0.001) via meta-analysis with larger WES (n=20,316) and array genotype datasets (n=19,033) from these same cohorts, using independent samples (n_WES_=15,205, n_array_=15,125) not included in the original WGS experiment (see **Methods**)^14^. We developed a strategy to genotype coding CNVs from WES data using read-depth information from XHMM^28^, and measured copy number at the 20,058 exons intersecting with 819 candidate CNVs from WGS. We found that 281 exons from 392 CNV calls were able to recapture the copy number variability detected by WGS (at R^2^>0.1). To genotype SVs using array data, we used standard imputation methods to impute 2,127 bi-allelic SVs based on the background of array-genotyped SNPs (see **Methods**). The estimated imputation accuracy of SVs corresponding well to their LD with nearby SNPs, as expected (**Figure S3**). To assess performance more rigorously for the eight significant SVs described below, we also performed a leave-one-out experiment, and the validation rate ranged from 93.3%-99.8% (**Table S4**). Overall, we were able to accurately genotype 2,053 of 4,864 candidate SVs using exome (n=392) and/or array genotype data (n=1,705). We then ran single-variant tests on those genotyped SVs with the corresponding candidate traits in the independent samples, and performed a meta-analysis to calculate a combined p-value (**Table 2**).

After merging fragmented SVs, we ended up with 15 independent loci associated with 31 traits at genome-wide significance, 9 of which remained significant after correction for the multiple phenotypes. **Table 2** shows the summary statistics of the lead SVs for their top traits (see also **Table S5** for pre-merged summary statistics).

### Deletion of the *ALB* gene promoter is associated with multiple traits

The strongest signal in the combined study was a 4kb deletion immediately upstream of the *ALB* gene, affecting the promoter region (**Figure 3**). This variant was 16-fold enriched in the Finnish population compared to non-Finnish Europeans from 1KG (MAF: 1.6% vs. 0.1%) and was associated with 16 traits at genome-wide significance (**Table S5, Figure S4**). The top two associations were with serum albumin (p=1.47×10^−54^) and total cholesterol (p=1.22×10^−28^), and these are independent signals based on conditional analyses. The cholesterol signal appears to explain the remaining 14 trait associations, all of which are highly correlated (**Figure S4**). This SV was well-tagged by nearby SNPs (R^2^=0.73), and the tagging SNPs showed similar trait association patterns. To tease apart potentially indirect associations caused by LD, we performed fine-mapping analysis for serum albumin and total cholesterol with CAVIAR^32^ including the deletion variant and the 100 most significant SNPs on chr4:67-79Mb (see **Methods**). The top candidate for the association with total cholesterol was a SNP (rs182695896) in moderate LD (R^2^=0.49) with the deletion. Accounting for this SNP via conditional analysis attenuated the association between the deletion and total cholesterol (p=0.023, n=4014). The deletion was identified as the most probable causal variant for the association with albumin, and the association between the deletion and albumin remained significant after adjusting for rs182695896 (p=6.52×10^−13^, n=3,117). We also observed different causality patterns for the two traits by aligning the posterior probabilities with the LD structure of the causal candidates in 95% confidence sets (**Figure 3**). Thus, we hypothesize that the promoter deletion directly affects serum albumin by altering *ALB* gene expression, and is associated with total cholesterol through its genetic correlation with other underlying causal variant(s) in the same LD block.

**Figure 3.**
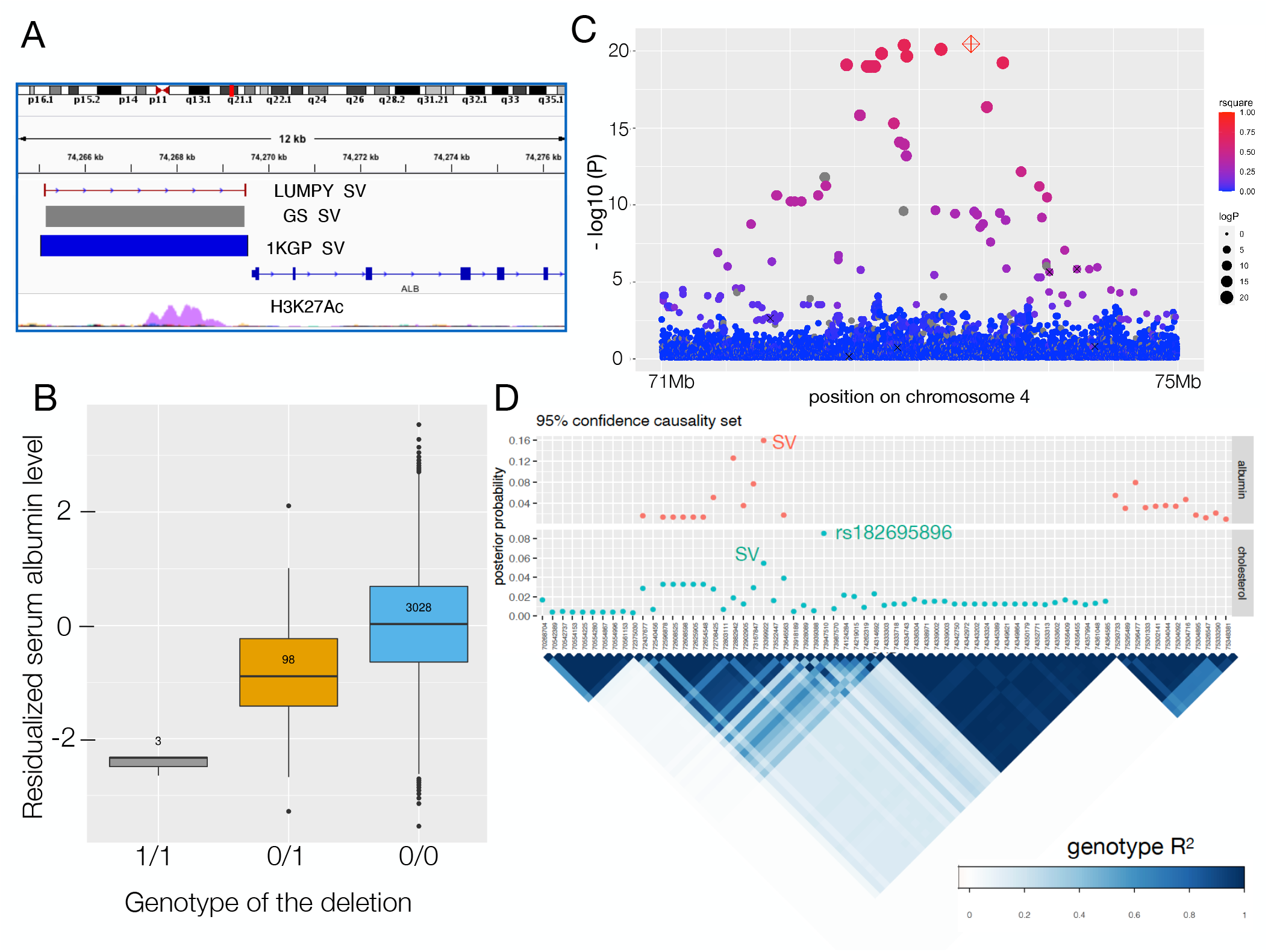
The *ALB* promotor deletion associated with serum albumin level and cholesterol traits. **(A)** The genomic location of the chr4 deletion, with coordinates detected from LUMPY, GenomeSTRiP and 1KG. The H3K27Ac track is from the ENCODE ^56^ data obtained from the UCSC genome browser. **(B)** Boxplot showing serum albumin levels stratified by genotype, with the sample size of each genotype group annotated at the center of each box. The trait value on the y-axis is the inverse normalized residual of raw measurement (residualized for age, age^2^, and sex). **(C)** Local Manhattan plot of albumin association signals on chr4:71-75Mb, including the *ALB* deletion (red diamond) and SNPs with minimum allele count of 9 (filled circles). The sizes of the circles are proportional to −log10(p) and colors indicated LD (Pearson R^2^) with the deletion (NA shown in grey). Six of the seven previously published GWAS signals are indicated with ‘x’ (the seventh was too rare in our data to be included in the test). **(D)** Fine-mapping results at the *ALB* locus for albumin and total cholesterol trait associations, using CAVIAR. The top panel shows the 95% confidence causality sets for albumin (top) and cholesterol (bottom) and posterior probability of each variant to be causal (assuming a maximum of two causal variants). The bottom panel shows the LD structure for the candidate variants, using the genotype correlation (Pearson R^2^) calculated from WGS data.

Prior studies^35–38^ have reported five albumin associated SNPs and two cholesterol associated SNPs in this region. In our conditional analyses including all intrachromosomal GWAS hits^39^, the SV-albumin association remained genome-wide significant (**Table 2**) while the SV-cholesterol association was diminished (conditioned p=0.004). To investigate the relationship between our signal and each of the seven previous GWAS SNPs, we tested the SV for association while conditioning on the reported SNPs one at a time (**Table S6**) and ran the association tests on those SNPs with the SV as covariate (**Table S7**). These results suggest that the *ALB* deletion is the causal variant for three prior albumin associations (rs16850360, rs2168889, and rs1851024), is linked to one previously reported cholesterol association (rs182616603), and is independent of two prior albumin associations (rs115136538, rs184650103) and one cholesterol association (rs117087731).

We next explored the potential downstream effects of this promoter deletion in the FinnGen dataset^40^, which reports GWAS results for 1,801 disease endpoints in 135,638 individuals. We queried the top SV-tagging SNP (rs187918276, R^2^=0.73) in the PheWeb browser^40^ (**Figure S5**); the top association was with statin medication use (p=6.5×10^−69^). The second set of signals appeared in the “Endocrine, nutritional and metabolic diseases” category, led by disorders of lipoprotein metabolism and other lipidemias (p=1.4×10^−11^), pure hypercholesterolemia (p=3.0×10^−11^), and metabolic disorders (p=1.8×10^−7^). These results support the medical relevance of genetic variation at this locus suggested by this and prior work; however, it is unclear whether these results are due to the *ALB* promoter deletion or the linked variants (e.g., rs182695896) associated with cholesterol.

### A multi-allelic CNV at *PDPR* is associated with pyruvate and alanine levels

We identified a cluster of 13 highly correlated CNV calls at chr16q22.1 that were strongly associated with pyruvate (p=4.81×10^−21^) and alanine (p=6.14×10^−12^) levels in the serum. We reconstructed the copy number profile of this locus from short-read WGS data (see **Methods**) and confirmed that the 13 correlated variant calls correspond to a single ∼250kb multiallelic CNV (CNV1 in **Figure 4**) spanning the coding sequence and 5’ region of *PDPR*, a gene involved in the pyruvate metabolism pathway. *PDPR* encodes the regulatory subunit of pyruvate dehydrogenase phosphatase (PDP) which catalyzes the dephosphorylation and reactivation of pyruvate dehydrogenase complex, the catalyst of pyruvate decarboxylation. According to this mechanism, fewer copies of *PDPR* should slow down the decarboxylation reaction and lead to increased pyruvate levels, and increased copies should decrease pyruvate levels, consistent with our data (**Figure 4**). This CNV was also negatively associated with alanine levels, the product of pyruvate transamination, and conditional analysis suggested this association was mediated through pyruvate (**Table S8**).

**Figure 4.**
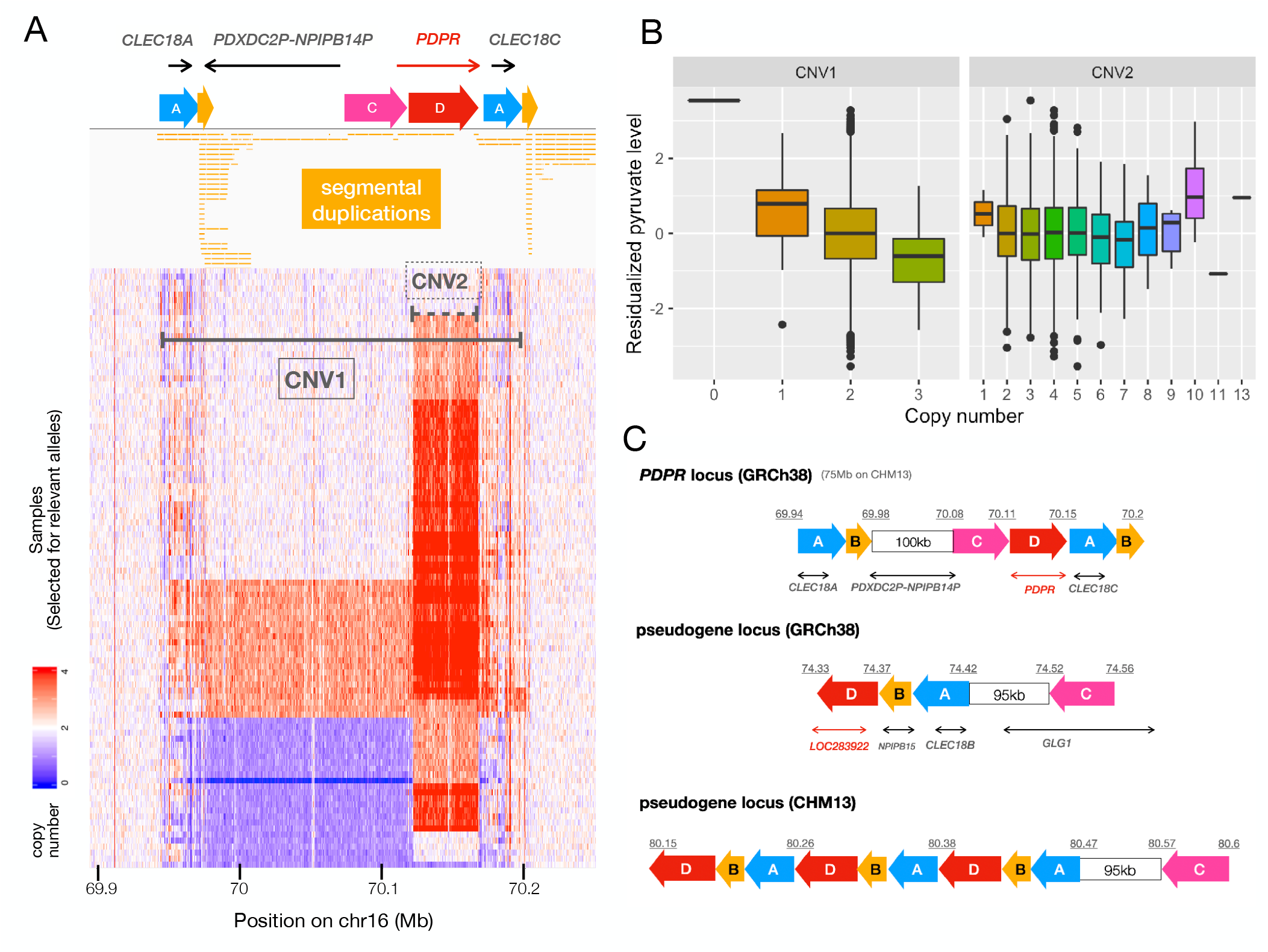
The multi-allelic CNV at the *PDPR* locus affecting pyruvate and alanine. **(A)** The *PDPR* locus showing (from top to bottom) genes, duplicated genomic segments based on dotplot analysis (see **Figure S6**), segmental duplication annotations from the UCSC table browser^57^, and copy number profiles for 100 samples comprising 51 carriers and 49 non-carriers for CNV1. Copy number is shown in 500bp windows, as determined by CNVnator, and the color saturates at four copies. The two horizontal lines indicate locations of the two CNVs (solid-CNV1, dashed-CNV2). **(B)** Pyruvate levels for 3,121 WGS samples stratified by copy number genotypes of CNV1 (p=9.41×10^−11^) and CNV2 (p=0.6). **(C)** Structure of GRCh38 reference and CHM13 assembly at the *PDPR* locus (top) and its pseudogene locus (bottom two), using the same annotations as in part (A). Blocks with the same color and letter notation are highly similar DNA sequences and arrows show the direction of alignments. Diagrams were drawn based on the dot plots in **Figure S6**. The segment B corresponds to LCR16a, the core element shared by many duplicons sparsely distributed on chromosome 16^41^.

An intriguing aspect of the *PDPR* locus is that it contains numerous segmental duplications (SDs), including highly similar local SDs scattered throughout the *PDPR* locus, additional SDs at a *PDPR* pseudogene (*LOC283922)* located 4 Mb distal to *PDPR*, as well as more divergent copies located ∼55Mb away on chr16p13.11. These include LCR16a, a core element shared by many SDs on Chr16 and a well-known driver of the formation of complex segmental duplication blocks in the genomes of humans and primates^41–43^. There are both duplication and deletion alleles of the *PDPR* gene, and these have indistinguishable breakpoints that correspond to LCR16a duplicons, suggesting these CNVs were caused by recurrent non-allelic homologous recombination. Similar to the *ALB* deletion described above (and many prior coding associations^14^), this CNV appears to be enriched in the Finnish population: the duplication allele was identified in 1KG with a frequency of 0.005 in non-Finnish Europeans, 50x less than the 0.025 frequency observed in our Finnish sample, and the deletion allele was not detected in 1KG. The CNV is poorly tagged by flanking SNPs (max R^2^<0.088), making it virtually undetectable using standard GWAS methods.

In addition, a second highly polymorphic and multiallelic CNV (CNV2 in **Figure 4**) intersects with CNV1 and covers >90% of the gene body of *PDPR*, missing the first three exons. Notably, CNV2 did not show association with pyruvate levels in our data (p=0.6), despite being previously reported as a *cis*-eQTL for *PDPR* in multiple tissues^10^. To resolve the structure of this locus, we aligned chromosome 16 of the GRCh38 reference against itself and also against the recent high-quality CHM13 assembly^44^ created from long-read sequencing data (**Figure S6**). Interestingly, we found that the sequence of CNV2 contains three inverted paralogs of the *LOC283922* locus (a *PDPR* pseudogene) in the CHM13 assembly, while there is only one copy of *LOC283922* in GRCh38 (**Figure 4**). These data suggest that CNV2 reflects highly variable structural alleles of *LOC283922* located 4Mb away from *PDPR*, and thus it is not surprising that this CNV does not affect pyruvate levels.

### Additional trait-association signals

We confirmed a previously reported association between the recurrent *HP* deletion and decreased total serum cholesterol levels^4^. In our data, this same deletion was strongly associated with serum glycoprotein acetyls quantified by NMR (p=3.53×10^−35^), and conditional analysis showed that the two associations were independent (**Table S8**). Since Boettger et al.^4^ proposed a plausible mechanism for the association of *HP* copy number and cholesterol, here we focus on the glycoprotein association. As a serum glycoprotein, haptoglobin forms dimers in individuals with the HP1/HP1 genotype (homozygous deletion) but forms multimers in individuals carrying HP2 allele(s). The multimers can be as large as 900kDa – more than twice the size of the dimers (86kDa)^45^ – which could result in fewer haptoglobin molecules in HP2 carriers, and consequently fewer glycoprotein molecules overall.

We identified five trait associations involving common SVs that were within 1Mb of previously published GWAS loci for the same traits. All SVs were well-tagged by SNPs (R^2^>0.9) and were either intronic or upstream of genes that are functionally related to the associated phenotypes. In all five cases there were stronger SNP signals nearby, and the SV associations dropped to not more than nominal significance when conditioned on the known GWAS SNPs (**Table 2**). This suggests that instead of having independent effects on the phenotypes, those SVs were more likely to be in LD with the causal variants.

Additionally, we identified a low-frequency (MAF=0.01) SV associated with serum tyrosine levels (combined p=4.17×10^−10^). This variant was a 4kb deletion of *IL34*, affecting the first exon of one transcript isoform and the intronic region of the two longer isoforms. There is a stronger signal from a SNP (rs190782607, p=1.44×10^−11^) within 100kb of and partially tagging the SV (R^2^=0.61), indicating that the SV is unlikely to be the causal variant. However, the p-value of this association remained at a similar level when conditioned on known GWAS SNPs^39^ (**Table 2**), suggesting a novel signal. *IL34* mediates the differentiation of monocytes and macrophages and to our knowledge has not previously been reported to be associated with amino acid traits^46^. *IL34 is* a crucial gene in the immune pathway and one study^47^ reported altered phenylalanine to tyrosine ratios associated with the immune activation and inflammation in CVD patients, which could explain the initial association as immune response related amino acid change. In addition, several studies^48,49^ have reported increased serum *IL34* levels in some cardiometabolic diseases that could potentially serve as a biomarker^50,51^.

The re-discovery of known loci described above demonstrates the effectiveness of our study design. Our CNV detection pipeline also detected two associations with metabolic traits that appear to be related to blood cell-type composition rather than inherited genetic variation. We identified three clusters of CNVs on chr7q34, chr7p14 and chr14q11.2 associated with C-reactive Protein (CRP) levels in the plasma, a biomarker for inflammation and a risk factor for heart disease (**Table 2, Table S5**). These CNVs are large, involve subtle alterations in copy number, and correspond to T cell receptor loci, suggesting that they are likely to reflect somatic deletions due to V(D)J recombination events during T cell maturation. This hypothesis was supported by the read-depth coverage pattern (see **Figure S7)**, where the measured copy number is lowest at the recombination signal sequence (RSS) used constitutively for rearrangement, and gradually increases with increasing distance to the RSS. The cause of this association is unclear but may reflect increased T-cell abundance and CRP levels due to active immune response in a subset of individuals. Interestingly, the CNVs were also associated with serum NMR tyrosine and serum NMR histidine (**Table S5**), which potentially supports the findings of previous publications about the involvement of amino acid metabolism in immune response^52,53^.

Interestingly, we also indirectly measured mitochondrial (MT) genome copy number variation due to the mis-mapping of reads from mitochondrial DNA to ancient nuclear MT genome insertions (NUMT)^54^ on chromosomes 1 and 17, that show strong homology to segments of the MT genome. These apparent “CNVs”, which reflect MT abundance in leukocytes, were strongly associated with fasting insulin levels (p=1.00×10^−10^) and related traits, and are the topic of a separate study^55^.

We also discovered three association signals corresponding to dense clusters of fragmented CNV calls within highly repetitive and low-complexity regions including simple repeats and segmental duplications (**Table 2**). Interpreting patterns of variation and trait association at these loci remains challenging due to their complex and repetitive genomic architecture, and known alignment artifacts within such regions. Although we were not able to identify any technical artifacts that might explain these specific associations, they should be interpreted with caution. Further investigation of these highly repetitive loci will require improved sequencing and variant detection methods.

## Discussion

We have conducted what is to our knowledge the first complex trait association study based on direct ascertainment of SV from deep WGS data. Our study leverages sensitive SV detection methods, extensive cardiometabolic quantitative trait measurements, and the unique population history of Finland. Despite the relatively modest sample size and limited power of this study, we identified 9 novel and 6 known trait associated loci. Most notably, we identified two novel loci where SVs are the likely causal variants and have strong effects on disease-relevant traits. Both SVs are ultra-rare in non-Finnish Europeans but present at elevated allele frequency in Finns – presumably due to historical population bottlenecks and expansions – which mirrors the findings from our recent study of coding variation, where many cardiometabolic trait-associated variants were enriched in Finns^14^. The first, a deletion of the *ALB* promoter, strongly decreased serum albumin levels in carriers (∼1 standard deviation per copy), and also resides on a haplotype associated with cholesterol levels. This example shows that non-coding SVs can have extremely large effects, consistent with our prior results based on eQTLs^10^ and selective constraint^18^, and points to the importance of including diverse variant classes in trait association efforts . Although more work is required to understand the disease relevance of this deletion variant, we note that low levels of albumin can cause analbuminemia, which is associated with mild edema, hypotension, fatigue, lower body lipodystrophy, and hyperlipidemia.

The second, a multi-allelic CNV with both duplication and deletion alleles that affect *PDPR* gene dosage, has strong effects on pyruvate and alanine levels. Notably, this CNV is the product of recurrent NAHR between flanking repeats at a complex locus that has accumulated numerous segmental duplications over evolutionary time, and is not well-tagged by SNVs. This phenomenon – recurrent CNVs at segmentally duplicated loci – has been studied extensively in the context of human genomic disorders and primate genome evolution, but there are few examples for complex traits. This result underscores the importance of comprehensive variant ascertainment in WGS-based studies of common disease and other complex traits. We further note that it is unusual to observe multiallelic CNVs at a conserved metabolic gene such as *PDPR*; it is tempting to speculate about the role of such variation in human evolution.

Interestingly, our study also identified two novel and highly atypical trait associations that appear to be caused by variable cell type composition in the peripheral blood. Identifying these results was only possible due to our use of WGS on blood-derived DNA, combined with sensitive SV analysis methods capable of detecting sub-clonal DNA copy number differences. Our quantitative detection of subclonal T-cell receptor locus deletions formed by V(D)J recombination served as a proxy for measuring T cell abundance, and led to the novel result that CRP levels are associated with T cell abundance. We hypothesize that this association is caused by active immune response in a subset of individuals. Similarly, our quantitative detection of mitochondrial genome copy number via apparent “CNVs” at NUMT sites in the nuclear genome led to the novel and important finding that variable abundance of neutrophils vs. platelets in peripheral blood is strongly associated with insulin, fat mass, and related metabolic traits (as described in detail elsewhere^55^).

Taken together, these results highlight the potential role of rare, large-effect SVs in the genetics of cardiometabolic traits, and suggest that future comprehensive and well-powered WGS-based studies have the potential to contribute greatly to our understanding of common disease genetics.

## Data Availability

METSIM WGS, METSIM WES, and FINRISK WES sequence data are available through dbGaP (accessions phs001579, phs000752, and phs000756). METSIM variant and phenotype data will soon be available through AnVIL (accessions TBD). Genomic and phenotypic data for the FINRISK cohort are or will soon be obtainable through THL Biobank, the Finnish Institute for Health and Welfare, Finland (https://thl.fi/en/web/thl-biobank). Structural variant site frequency information is available in dbVAR (accession TBD). Summary statistics are available on GitHub (see **Web Resources**).

## Description of Supplemental Data

Supplemental Data include seven figures and eight tables.

## Acknowledgements

We thank D. Ray from Johns Hopkins University for her comments to the manuscript. This work was funded by an NHGRI CCDG award to IMH and NOS (UM1 HG008853) and DK U01 DK062370, the NHGRI large-scale sequencing grant (grant number 5U54HG003079), the Sigrid Jusélius Foundation (to SR), the University of Helsinki HiLIFE Fellow grants 2017-2020 (to SR), the Academy of Finland Center of Excellence in Complex Disease Genetics (grant number 312062 to SR, grant number 312074 to SR and AP), the Academy of Finland (grant number 285380 to SR), the National Heart, Lung and Blood Institute (grant number T32HL007081 to EY), and the National Center for Advancing Translational Sciences (grant number UL1TR002345 to EY). The funders had no role in study design, data collection and analysis, decision to publish, or preparation of the manuscript. We thank the MGI administration and data production team, in particular R. Fulton, L. Fulton, C. Fronick, A. Wollam, S.K. Dutcher, and J. Milbrandt. The FINRISK samples used for the research were obtained from THL Biobank. We thank all study participants for their generous participation in the THL Biobank, FINRISK study, and METSIM study. ASH was supported by the Academy of Finland (grant no. 321356). LC was supported by the McDonnell International Scholars Academy Fellowship. AJS was supported by the Mr. and Mrs. Spencer T. Olin Fellowship for Women in Graduate Study.

## Author Contributions

I.M.H. and N.O.S. conceived and directed the study. L.C., H.J.A, and I.D. adapted the GenomeSTRiP pipeline to perform CNV detection at scale. H.J.A. developed the pipeline for CNV genotyping based on CNVnator. L.C. and I.D. created the GenomeSTRiP callset; L.C. and H.J.A created the CNVnator callset; D.E.L. and K.L.K created the LUMPY callset, and led data management. L.C. led all analyses related to trait association, SV genotyping using WES and array data, and investigation of candidate loci. H.J.A, D.E.L, I.D, L.G. and A.A.R. led GATK callset creation and QC for WGS data. A.P., S.R., M.L, and J.K. contributed samples and phenotypic data. All authors edited the manuscript and/or provided intellectual contributions. L.C. and I.M.H. wrote the manuscript.

## Web Resources

The summary statistics of all the tested SVs and traits are available through GitHub: https://github.com/hall-lab/FinnSV_paper_1220

