## Supplemental figures and tables for "Association of Structural Variation with Cardiometabolic Traits in Finns"

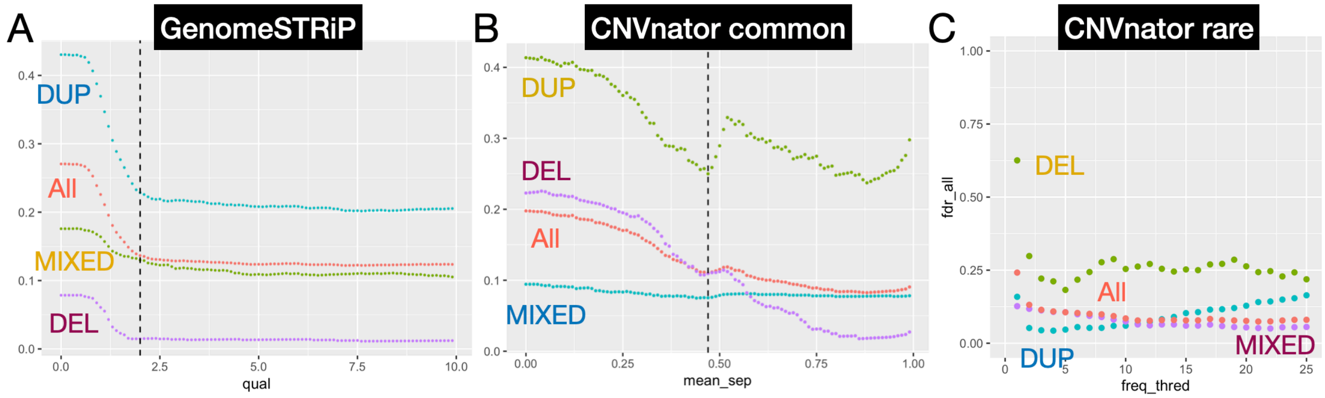


**Figure S1**. FDR curves under different quality thresholds for **(A)** GenomeSTRiP CNVs, **(B)** Common variants of CNVnator CNVs, and **(C)** Rare variants of CNVnator CNVs. The FDR was estimated from the array intensity data of METSIM samples using IntensityRankSumAnnotator from the GenomeSTRiP pipeline, among CNVs covered by at least two probes. GenomeSTRiP CNVs were filtered based on the "GSCNQUAL" score output by the software, common CNVnator CNVs were filtered by the "mean_sep" metrics from the constrained GMM model, and the rare CNVnator CNVs were filtered by carrier frequency. The results are presented for all variants as well as by different variant types, indicated by the colors shown.

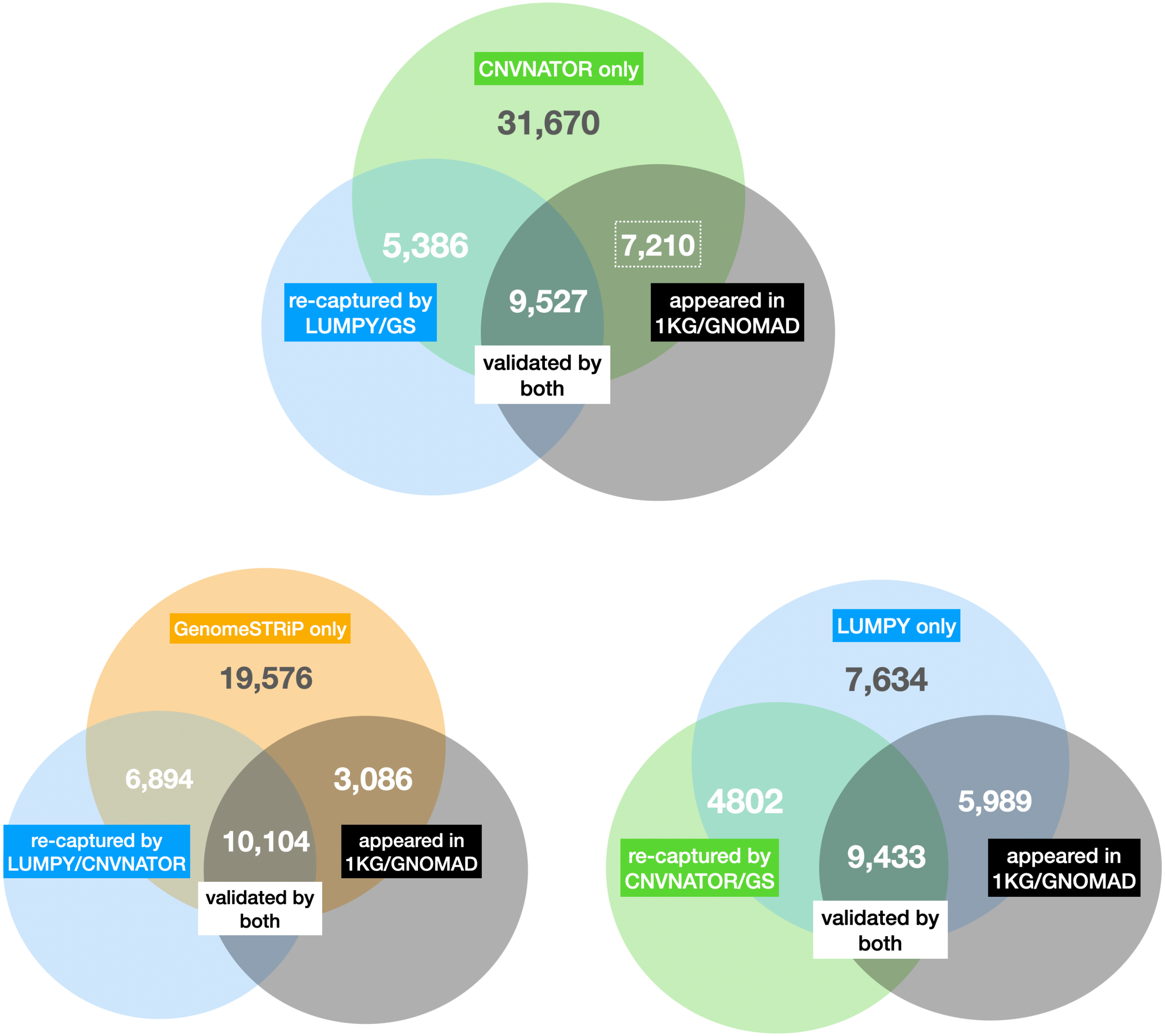


**Figure S2.** For each of the three SV detection methods used in this study, these venn diagrams show the number of CNVs that were also identified by the other two "internal" pipelines used in this study (left), and the "external" reference SV callsets from 1KG and gnomAD (right). The upper part of each diagram also shows the number of CNVs only identified by a given pipeline. Dashed rectangles were used to emphasize the number of CNVnator CNVs that were validated by external callsets but missed by the other two pipelines, showing the complementary nature of the methods used for this study. 50% reciprocal overlap was used to compare CNV calls from different callsets.

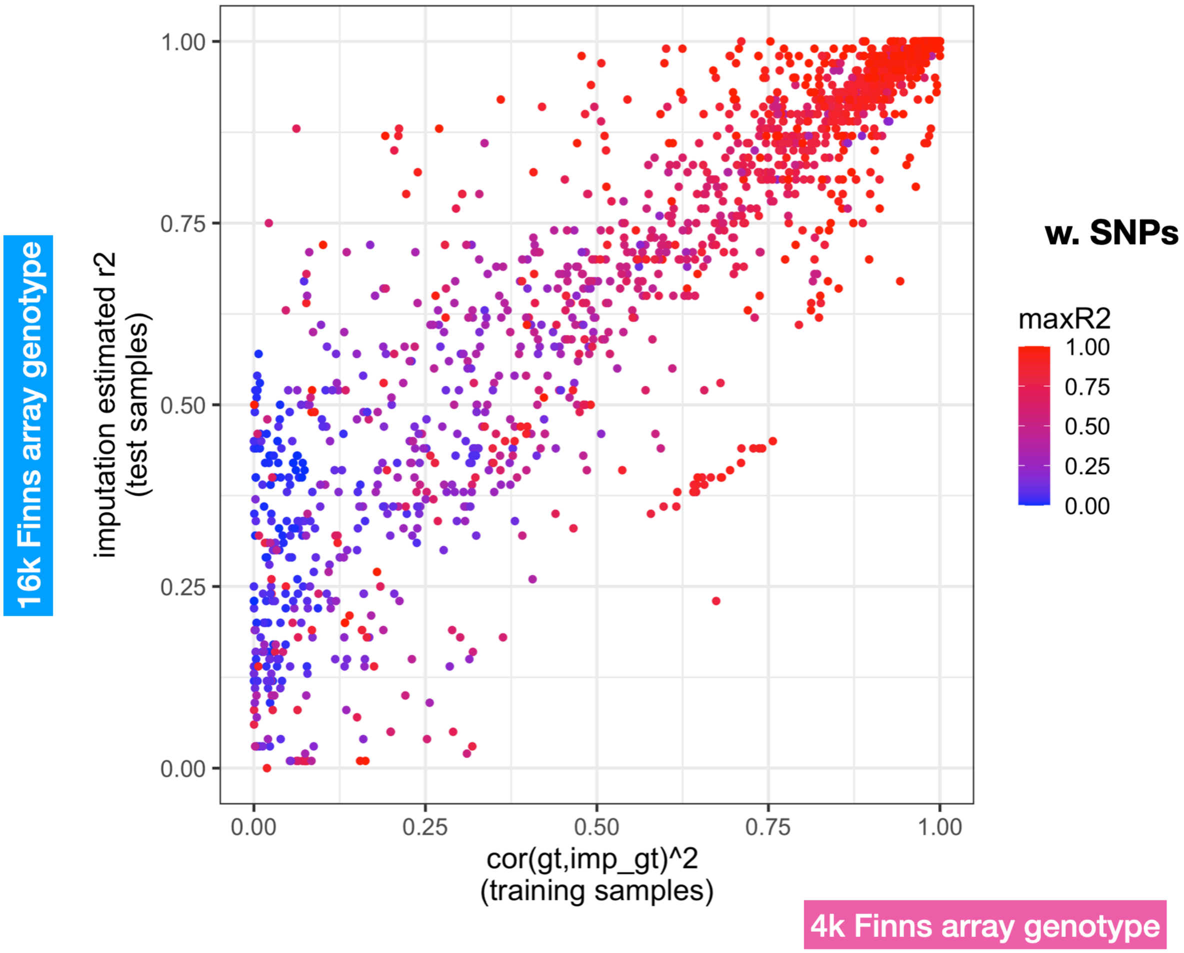


**Figure S3.** The overall evaluation of imputation quality with two metrics. The Y-axis shows the Beagle output quality score (DR2) for the ~15k tested samples, which is the estimated correlation between the imputed genotype and real genotype for each variant, and the X-axis shows the "training error" for the ~4k samples with WGS data. Training error was calculated using the WGS data as reference and array data as test input, after which the correlation of real genotype (based on WGS) and predicted genotype was calculated. The color shows how well each SV was tagged by nearby SNPs located within 1 Mb.

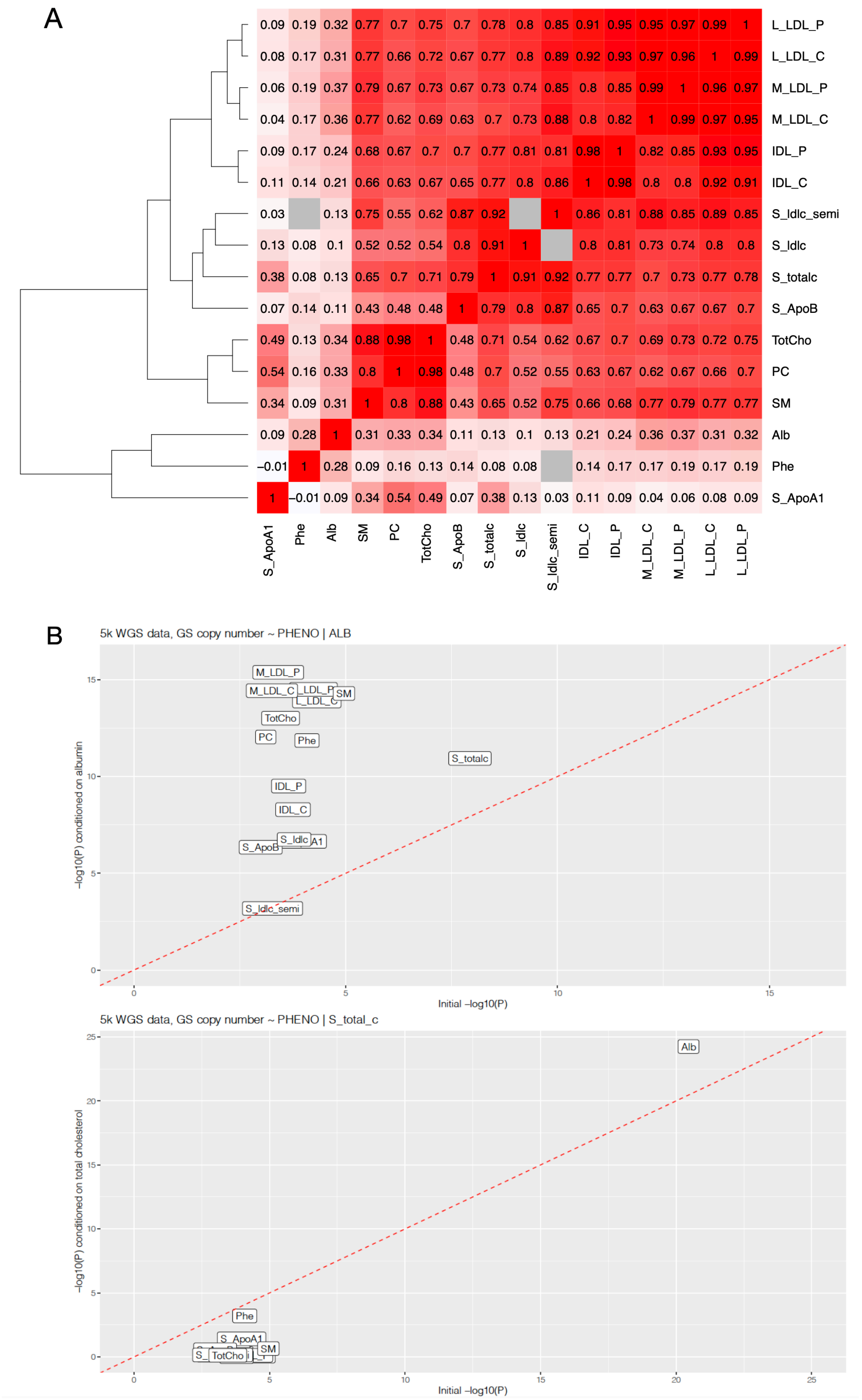


**Figure S4.** (**A)** The pairwise correlation (Pearson R) of the 16 traits that were significantly associated with *ALB* deletion. The cells shown in gray represent missing data, since the S_ldlc_semi trait (serum LDL cholesterol in semi-fasting samples) shared zero samples with S_ldlc (serum LDL cholesterol in fasting samples) and Phe (phenylalanine). **(B)** Comparison of the association p-value of the *ALB* deletion and the 16 traits, with (y-axis) and without (x-axis) albumin (top) and total cholesterol (bottom)

as a covariate. The increases of significant level of most traits when conditioned on albumin were likely due to Berkson's paradox[1].

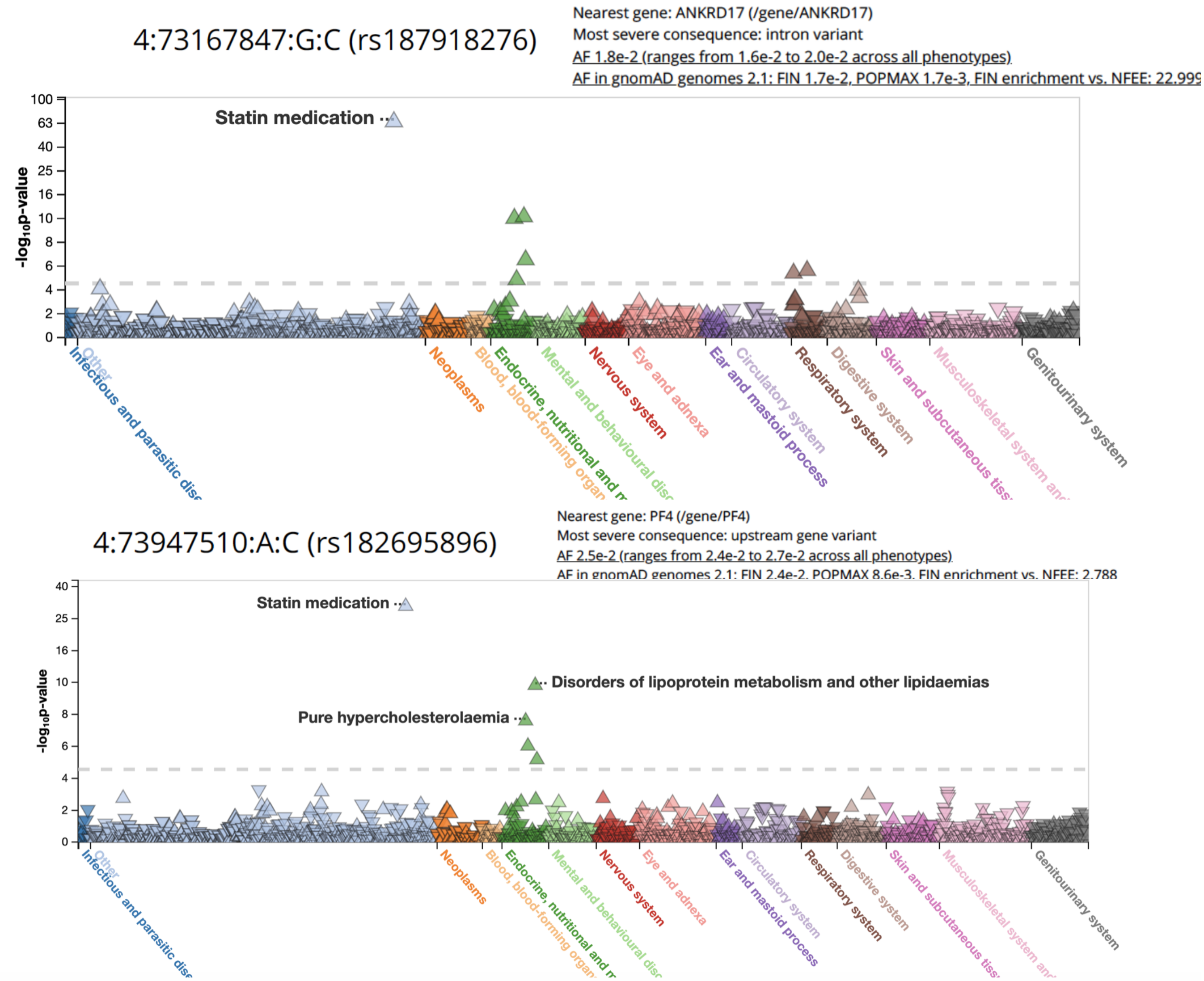


**Figure S5.** Screenshots from the FinnGen PheWeb browser[2] (Data Freeze 3) of the top tagging SNP for the *ALB* deletion (top) and for the cholesterol candidate (bottom) predicted by fine mapping with CAVIAR, showing the phenome-wide association results for each of the SNPs, colored by phenotype groups.

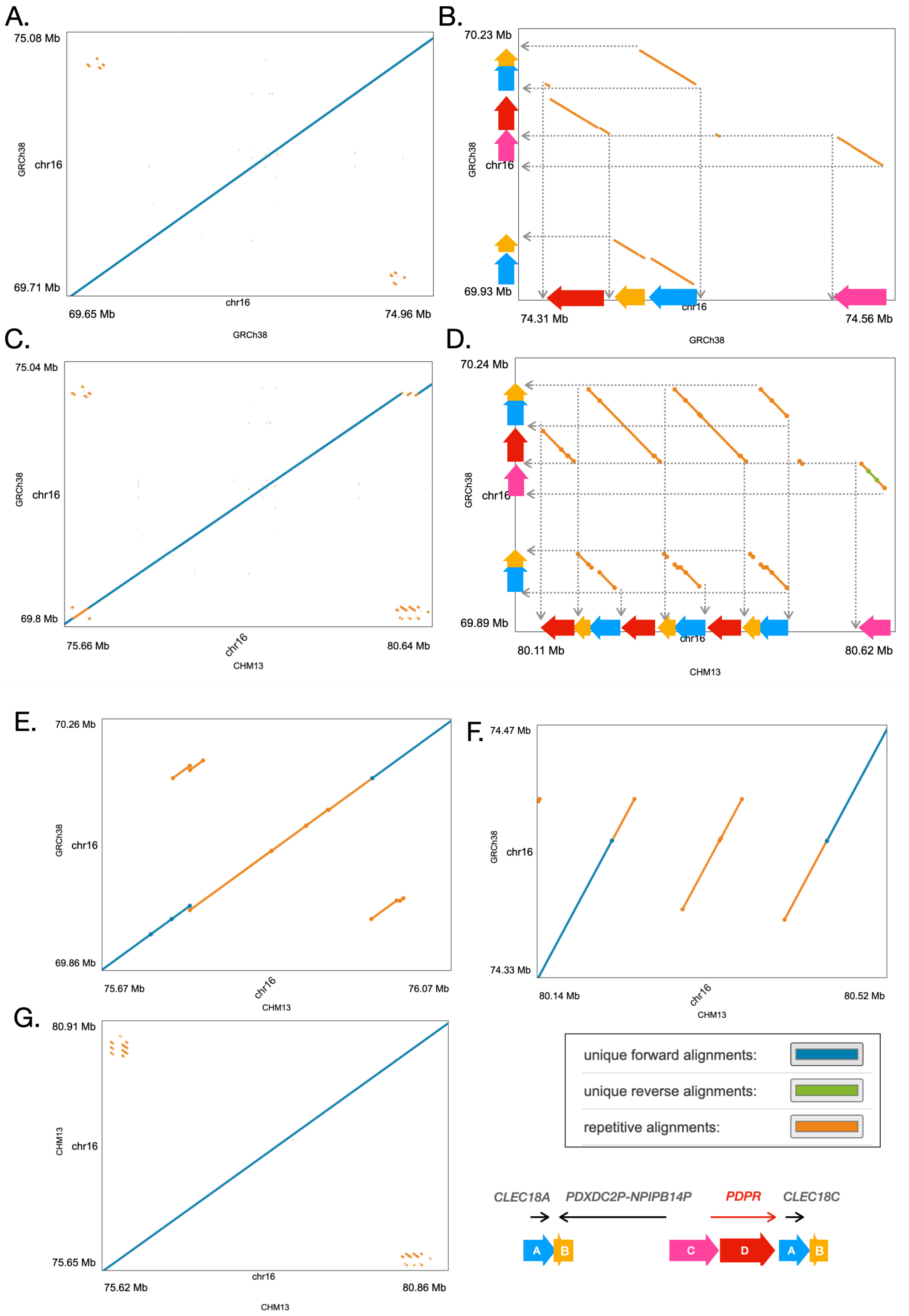


**Figure S6.** Dot plots showing the structure of the *PDPR* and nearby pseudogene locus in both the GRCh38 and CHM13 assemblies, with repetitive alignments shown in orange and unique alignments shown in blue and green (see legend bottom right). **(A)** The *PDPR* locus in GRCh38 (y-axis) aligned to the pseudogene locus (x-axis) in GRCh38, where **(B)** shows a zoomed-in version with the diagram used for **Figure 4** using the same colors and letter. **(C)** and **(D)** show the *PDPR* locus in GRCh38 vs.

the *PDPR* locus in CHM13. **(E)** and **(F)** show the pseudogene locus in GRCh38 vs. CHM13, and **(g)** shows the *PDPR* locus in CHM13 vs. itself.

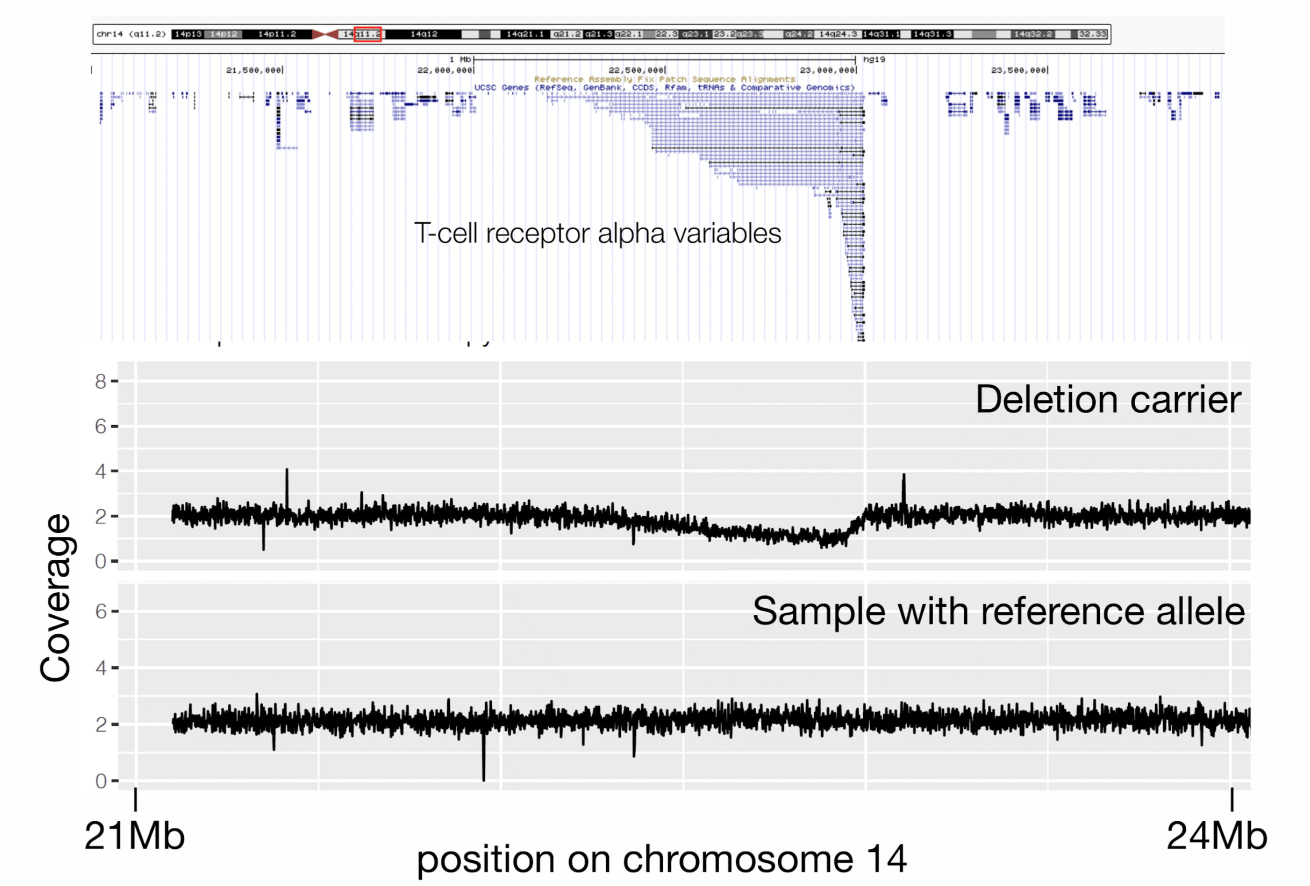


**Figure S7.** Read-depth coverage patterns at the chr14 T-cell receptor alpha variable region (coordinates LiftOver to GRCh37/hg19), showing one example for “deletion” carriers and one for a sample with the reference allele. The coverage values were calculated by CNVnator for 100bp windows, and the top gene track was extracted from UCSC genome browser (GRCh37/hg19).

### Supplemental Tables

**Table S1**. Variant and sample counts in each QC step for WGS data.

| | LUMPY | | GS | | CNVNATOR | | ALL_variants |
|---|---|---|---|---|---|---|---|
| | # variants | # samples | # variants | # samples | # variants | # samples | # variants |
| **Pipeline output** | 120793 | 5065 | 111141 | 5087 | 92862 | 4979 | 324796 |
| **Score-filtered** | 39392 | 5065 | 46702 | 5087 | 55371 | 4979 | 141465 |
| **FD-sites-filtered** | 39075 | 5065 | 45963 | 5087 | 54252 | 4979 | 139290 |
| **outlier-sample-filtered** | 39075 | 5062 | 45963 | 4966 | 54252 | 4967 | 139290 |
| **final-high-quality** | 37268 | 4848 | 43525 | 4848 | 53793 | 4848 | 134586 |
| **high-quality-autosome** | 35713 | 4848 | 39660 | 4848 | 53793 | 4848 | 129166 |
| **tested (MAC>9)** | 11633 | 4030 | 11062 | 4030 | 41877 | 4030 | 64572 |

**Table S1**. Variant and sample counts in each QC step for WGS data separated by variant calling pipelines. FD – false discovery, see **Methods** for the filtering criteria in each step.

**Table S2**. Genotype redundancy estimation

| Variants | CNVNATOR | LUMPY | GS | ALL | Tested_all |
|---|---|---|---|---|---|
| **Original count** | 53,793 | 35,713 | 39,660 | 129,166 | 64,572 |
| **VeffLi independent count*** | 24,330 | 27,676 | 29,445 | 71,688 | 26,495 |
| **Ratio** | 45.23% | 77.50% | 74.24% | 55.50% | 41.03% |
| **Genome-wide significant threshold** | - | - | - | - | 1.89E-06 |
| **Experiment-wide significant threshold**** | - | - | - | - | 3.32E-08 |
| * VeffLi results: sum of per chromosome estimates | | | | | |
| ** effective number of traits 56.8566 (ori:116) | | | | | |

**Table S2.** Estimation of redundant SV calls based on genotype information. Redundant variant calls identified by multiple SV detection methods are expected to have genotypes that are highly correlated. We therefore applied matSpDlite to each pipeline and to the combined callset to calculate the numbers of independent makers (VeffLi). We then applied the same method to the subset of the variants included in the trait association test and to the phenotypes to perform Bonferroni correction for the genome-wide significance threshold and experiment-wide threshold.

**Table S3.** Fragmentation level

| Pipeline | #. SV | # Cluster | average cluster size | % single variant cluster | size of the largest cluster |
|---|---|---|---|---|---|
| GS | 39,660 | 24,497 | 1.619 | 75% | 96 |
| LUMPY | 35,713 | 23,751 | 1.321 | 90% | 458* |
| LUMPY CNV | 27,858 | 21,759 | 1.28 | 90% | 149 |
| CNVNATOR | 53,793 | 16,962 | 3.171 | 73% | 527 |
| * a large inversion on chr7 with size of 44mb covered 400+ other variants | | | | | |

**Table S3.** Estimation of SV fragmentation based on physical clustering. Due to coverage fluctuations, CNV calls detected by read-depth analysis are often fragmented into multiple adjacent CNV calls that in fact represent a single variant. To estimate the degree of fragmentation, we clustered high-confidence autosomal CNVs within 10bp of each other and calculated the average number of SVs per cluster (average cluster size), the percentage of single variant clusters, and the maximum number of variants per cluster (size of the largest cluster).

**Table S4.** Leave-one-out validation for genome-wide significant SVs

| VAR | FALSE | TRUE | AC_RATE |
|---|---|---|---|
| 40551 | 17 | 3891 | 0.996 |
| 52933 | 113 | 3795 | 0.971 |
| 61703 | 55 | 3853 | 0.986 |
| 62003 | 7 | 3901 | 0.998 |
| chr12_95946601_95947800 | 260 | 3648 | 0.933 |
| chr16_72057601_72058200 | 63 | 3845 | 0.984 |
| chr20_45906701_45907200 | 144 | 3764 | 0.963 |
| CNV_chr4_73399922_73404147 | 9 | 3899 | 0.998 |

**Table S4.** The “leave-one-out” validation experiment to assess imputation quality of the eight genome-wide significant SVs. For each variant, we ran 3,908 imputation experiments and in each we used one sample as the test genome and the other samples as the reference. The accuracy rate was calculated among all 3,908 tests.

**Table S5.** Summary statistics for all genome-wide significant signals, before manually clumping redundant variants into single calls. P-value (P), effect size (BETA), allele count (AC), allele frequency (AF) and sample size (N) are shown for whole genome (WGS), exome (WES) and imputed (IMP) data. The combined p-value (COMBINED_P) was calculated using Fisher’s method.

**Table S6**. Test ALB deletion conditioned on published GWAS SNPs

| Supplementary Table 6. Test ALB deletion conditioned on published GWAS SNPs | | | | | | | | | |
|---|---|---|---|---|---|---|---|---|---|
| rs ID | GWAS trait | First author, year | R2 w. SV | MAF(Finns) | MAF (Reported) | SV P value | | Beta (conditional analysis) | |
| | | | | | | albumin ~ SNP + SV | cholesterol ~ SNP + SV | albumin | cholesterol |
| rs16850360 | albumin | Kettunen et al, 2012 Inouye et al, 2012 | 0.3 | 0.025 | 0.03 | 8.10E-18 | 6.00E-04 | 1.1 | -0.39 |
| rs182616603 | cholesterol | Surakka et al, 2015 | 0.3 | 0.024 | 0.01 | 6.60E-17 | 4.00E-03 | 1.06 | -0.32 |
| rs2168889* | albumin | Inouye et al, 2012 | 0.12 | 0.049 | 0.05 | 6.40E-23 | 9.70E-05 | 1.05 | -0.37 |
| rs1851024 | albumin | Inouye et al, 2012 | 0.08 | 0.049 | 0.05 | 2.30E-19 | 3.30E-08 | 0.91 | -0.5 |
| rs117087731 | cholesterol | Surakka et al, 2015 | 6.00E-04 | 0.02 | 0.01 | 2.50E-21 | 1.70E-08 | 0.91 | -0.49 |
| rs115136538 | albumin | Kettunen et al, 2012 | 3.00E-05 | 0.005 | 0.02 | 2.80E-21 | 1.50E-08 | 0.91 | -0.49 |
| rs184650103 | albumin | Kettunen et al, 2016 | 3.00E-05 | 0.001 | 0.01 | 2.90E-21 | 1.60E-08 | 0.91 | -0.49 |
| SV ~ albumin P value = 3.49E-21, beta = 0.9107 | | | | | | | | | |
| SV ~ cholesterol P value = 1.17E-08, beta = -0.4929 | | | | | | | | | |
| rs2168889*: collider effect | | | | | | | | | |

**Table S6.** Conditional analysis of the three multi-trait associated variants, taking one trait as a covariate and testing the other. Additional traits were tested for the *ALB* deletion conditioned on albumin and total cholesterol, the results of which can be found in **Supplementary Figure 6b**. The covariate trait was defined as a mediator of the tested trait if the conditional p-value failed the genome-wide significance threshold ($1.89x10^{-6}$). MAF(Finns) – MAF in our data, MAF(Reported) – MAF reported in previous GWAS studies.

**Table S7**. Test the published GWAS SNPs w./w.o. SV as covariate

| Supplementary Table7. Test the published GWAS SNPs w./w.o. SV as covariate | | | | | | | | | | | |
|---|---|---|---|---|---|---|---|---|---|---|---|
| rs ID | GWAS trait | First author, year | R2 w. SV | MAF(Finns) | MAF (Reported) | SNP P value | | Beta (alb) | SNP P value | | Beta (chol) |
| | | | | | | albumin ~ SNP | albumin ~ SNP + SV | | cholesterol ~SNP | cholesterol ~SNP + SV | |
| rs16850360 | albumin | Kettunen et al, 2012 Inouye et al, 2012 | 0.3 | 0.025 | 0.03 | 2.30E-06 | 5.00E-02 | -/+ | 2.90E-06 | 1.80E-01 | +/+ |
| rs182616603 | cholesterol | Surakka et al, 2015 | 0.3 | 0.024 | 0.01 | 1.40E-06 | 8.00E-02 | -/+ | 5.30E-08 | 2.00E-02 | +/+ |
| rs2168889 | albumin | Inouye et al, 2012 | 0.12 | 0.049 | 0.05 | >0.05 | - | - | 4.40E-07 | 2.00E-03 | +/+ |
| rs1851024 | albumin | Inouye et al, 2012 | 0.08 | 0.049 | 0.05 | 2.20E-03 | 8.30E-01 | +/+ | >0.05 | - | - |
| rs117087731 | cholesterol | Surakka et al, 2015 | 6.00E-04 | 0.02 | 0.01 | >0.05 | - | - | >0.05 | - | - |
| rs115136538 | albumin | Kettunen et al, 2012 | 3.00E-05 | 0.005 | 0.02 | >0.05 | - | - | >0.05 | - | - |
| rs184650103* | albumin | Kettunen et al, 2016 | 3.00E-05 | 0.001 | 0.01 | NA | NA | NA | NA | NA | NA |
| *Note: rs184650103 was too rare to be included in the test, so the summary statistics were marked as "NA", to differentiate from "-", which marks non-significant SNPs. | | | | | | | | | | | |

**Table S7.** Association analysis between the *ALB* deletion and albumin/total cholesterol conditioned on the seven previously published GWAS SNPs one at a time, ordered by the genotype correlation with the SV. None of the seven SNPs diminishes the SV-albumin signal, while the first three SNPs attenuate the SV-cholesterol signal, suggesting that they might also be in LD with the underlying causal variants for cholesterol.

**Table S8**. Conditional analysis (phenotype - phenotype)

| Variant | Tested trait | Covariate trait | P WGS | P conditioned | BETA WGS | BETA conditioned | Mediator? |
|---|---|---|---|---|---|---|---|
| ALB deletion | Albumin | Total Cholesterol | 3.49E-21 | 5.74E-25 | 0.9107 | 0.9937 | N |
| | Total cholesterol | Albumin | 1.17E-08 | 1.16E-11 | -0.4929 | -0.6558 | N |
| PDPR mCNV | Pyruvate | Alanine | 9.41E-11 | 6.59E-05 | -0.5817 | -0.4344 | N |
| | Alanine | Pyruvate | 2.93E-07 | 1.47E-03 | -0.5744 | -0.3197 | Y |
| HP deletion | Glycoprotein | Total cholesterol | 1.51E-11 | 2.78E-15 | -0.2081 | -0.1988 | N |
| | Total cholesterol | Glycoprotein | 1.01E-05 | 2.62E-10 | 0.1466 | 0.1604 | N |

**Table S8.** The association tests between each of the seven previously published GWAS SNPs and serum albumin/total cholesterol, with and without the *ALB* deletion as a covariate (SNPs with p-value > 0.05 were not included in the conditional analysis, with "-" in the related fields). The "Beta" column shows the direction of effects of SNPs with/without the SV in the model. rs115136538, rs184650103 and rs117087731 did not show significant association with either trait in our dataset. The other SNPs showed signals with albumin or total cholesterol which became much less significant after conditioning on SV genotype. *Note: rs184650103 was too rare to be included in the test, so the summary statistics were marked as "NA", to differentiate from "-", which marks non-significant SNPs.

2. PheWeb.
